# Highly multiplexed targeted plasma proteomics quantifies several hundred blood proteins in serum from colorectal carcinoma patients

**DOI:** 10.1101/2022.04.01.486663

**Authors:** Antoine Lesur, François Bernardin, Eric Koncina, Elisabeth Letellier, Gary Kruppa, Schmit Pierre-Olivier, Gunnar Dittmar

## Abstract

The rapid analysis of human serum and plasma can provide deep insights into changes of the blood proteome in response to different patient treatments or diseases. Targeted proteomics techniques, like SRM and PRM, can be utilized to monitor proteins at high sensitivitym but so far were limited to smaller protein panels, which can be monitored in one experiment. The recently, on a Bruker tims-TOF pro mass spectrometer, developed parallel reaction monitoring-parallel accumulation − serial fragmentation (prm-PASEF) method expands the standard PRM method by using ion-mobility. The use of ion mobility as a fourth separation dimension increases the proteome coverage while reducing the length of the necessary chromatogeaphic separation. By combining an isotope-labeled reference standard, which covers 579 plasma proteins, we were able to quantify 565 proteins in plasma using prm-PASEF, with the least abundant protein being quantified at 7 amol. We continued the analysis by combining the isotype-labeled reference standard with dia-PASEF, which allowed the quantification of 549 proteins. Both methods were used to analyze 20 patient plasma samples from a colorectal cancer (CRC) cohort. The analysis identified 16 differentially regulated proteins between the CRC patient and control individual plasma samples. 15 of the 16 proteins showed a high correlation to the mRNA expression in CRC tumor samples, showing the technique’ s potential for the rapid identification of potential biomarkers in larger cohorts, abolishing the need for preselection of potential biomarker proteins.

**Graphical abstract:** 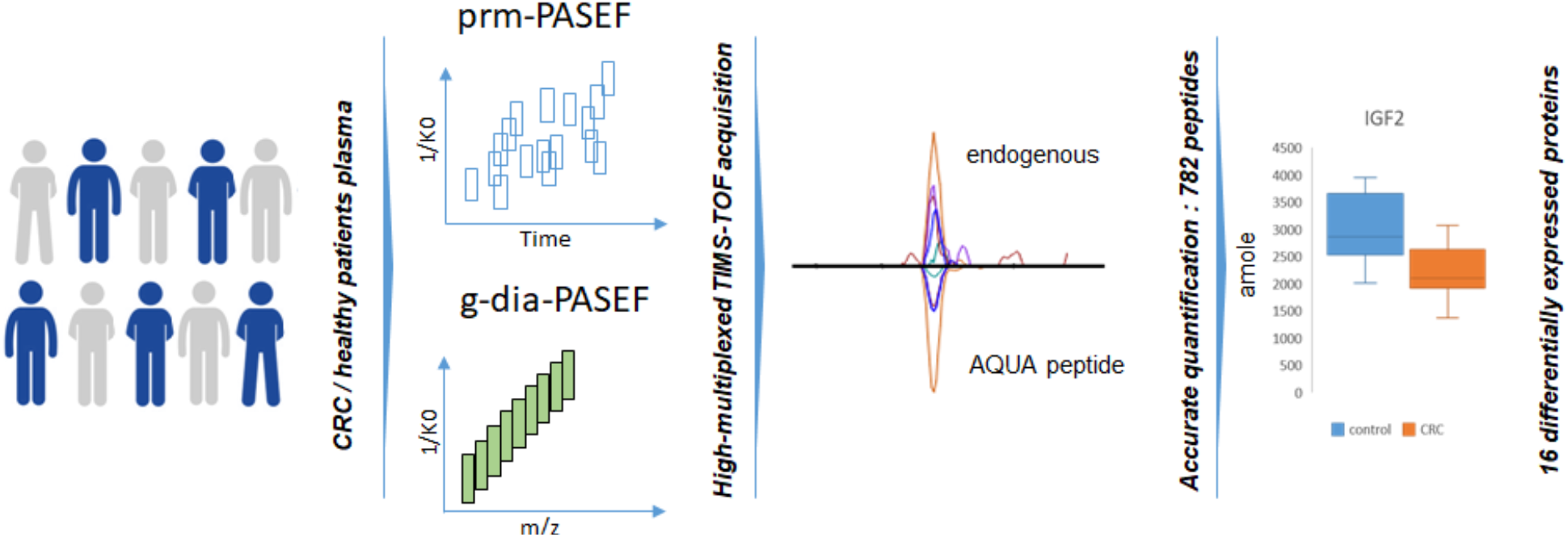

## Introduction

Blood-based diagnostics is one of the pillars for disease diagnosis at an early or asymptomatic stage, the prediction of the treatment outcome or a guide for the choice of the most appropriate treatment. Blood components including the serum and the plasma can be collected with minimal discomfort to patients and are readily available for prospective and retrospective studies. Although blood-based assays have been used for a long time, many diagnostics still rely on measuring a single protein or the composition of the cells contained within the blood sample.

Recently, it has been shown that protein panels provide a better diagnostic or predictive power than single protein markers^1–3^. As many described biomarkers are derived from the direct analysis of the affected tissue or tumor material, they usually do not directly translate into valid biomarkers in blood samples, making further studies and method development necessary ^4,5^.

As a multiplexed technique, mass spectrometry plays an integrated role in the discovery and verification of biomarkers candidates and is an alternative to antibody-based analysis techniques^6^.

The shotgun proteomic analysis of serum and plasma allows the identification of blood proteins which then serve as the basis for designing targeted mass spectrometry methods. Targeted proteomics provides reliable measurements of selected sets of proteins in large cohorts of samples without the problem of missed values associated with the shotgun proteomics. The detection limits are improved by focusing the MS on a limited set of targets.

The targeted mass spectrometric method spectrum includes SRM (selected reaction monitoring) using a triple quadrupole and the more recently developed PRM (parallel reaction monitoring) which takes advantage of the higher resolution and mass accuracy of an orbitrap or time-of-flight (TOF) mass analyzers. PRM increases the selectivity in complex matrices like plasma^7^ or serum and it has been successfully used to confirm proteins as biomarkers^7,8^.

We recently developed a new targeted acquisition method (prm-PASEF) that uses trapped ion mobility (TIMS) upfront a quadrupole time of flight mass spectrometer to increase the multiplexing capability without losing sensitivity^9^.

In this study, we employed prm-PASEF for the quantification of 565 proteins using 782 quantified isotope-labeled synthetic reference peptides (*i*.*e*. AQUA peptides). Additionally, we combined the use of the complex isotope-labeled reference peptides and combined it with dia-PASEF^10^. This version of dia-PASEF, called guided dia-PASEF (g-dia-PASEF), allows the rapid measurement of peptide samples with the option of absolute quantification (figure 1). The quantified reference peptides can be used to convert the MS signal of their endogenous counterparts into a concentration and thus allows the comparison of the quantification results across different mass spectrometry systems or acquisition methods. Both methods allowed measuring and quantifying a significant fraction of the blood proteome in a single assay. Using the plasma from 20 patients of a colon carcinoma cohort we showed the potential for the rapid identification of protein panels as potential disease markers.

**Figure 1.**
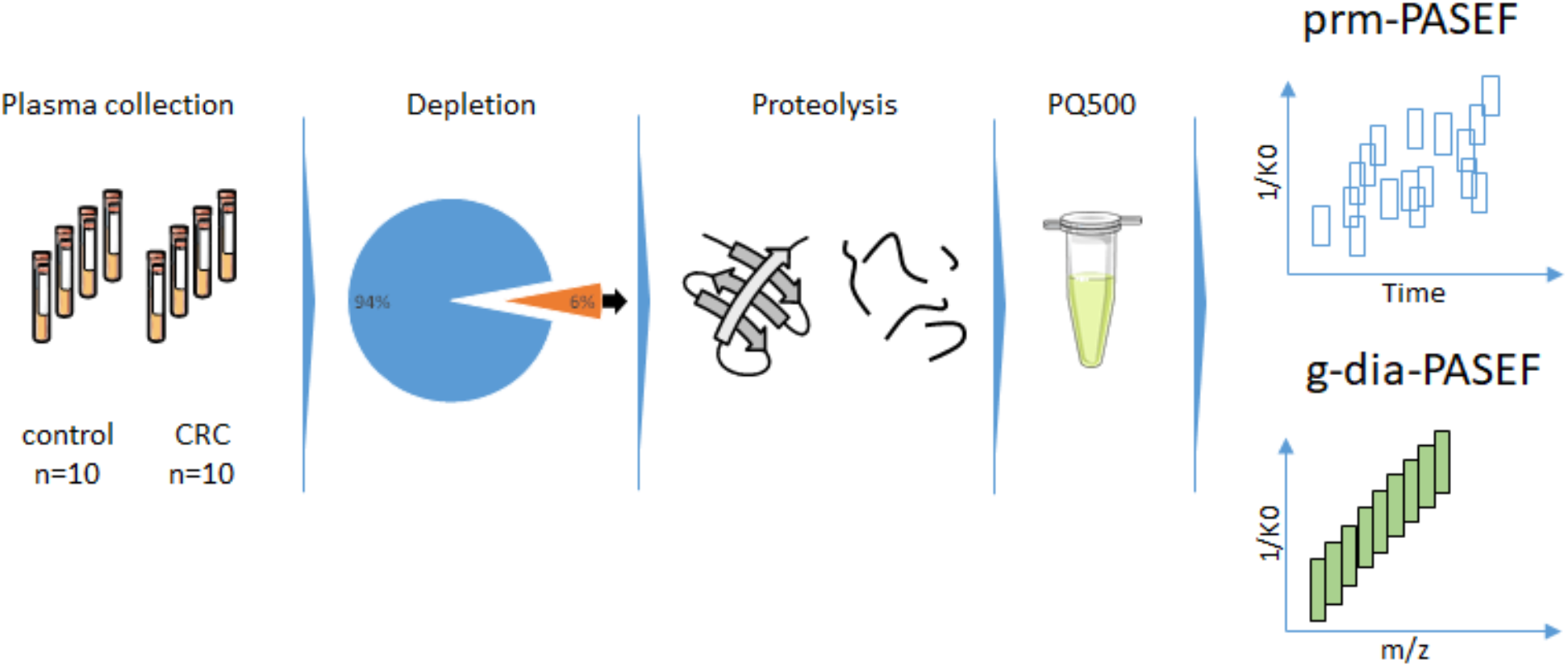
Plasma samples were depleted, digested by trypsin, and spiked with PQ500 isotope-labeled synthetic peptides. Samples were analyzed prm-PASEF and g-dia-PASEF, and data were processed by Skyline.

## Material and methods

### Plasma depletion and processing

Aliquots of 20 µL of human plasma were depleted on a 1260 Infinity Bio-inert LC system (Agilent) coupled to a depletion column (human 14 multiple affinity removal column; 4.6 × 50 mm; Agilent) according to the manufacturer’s procedure. After depletion, the buffer was exchanged to 100 mM NH_4_HCO_3_ and the volume was concentrated to 100 µL using a spin concentrator with a 5 kDa cutoff (Pall). Proteins were denatured with 1% sodium deoxycholate (SDC), reduced with 10 mM dithiothreitol for 30 min at 37°C, and alkylated with 25 mM iodoacetamide for 30 min at room temperature. All reagents were prepared in a freshly made 50 mM ammonium bicarbonate buffer. The denatured samples were diluted to 0.5% SDC and proteolysis was performed by the addition of 6,5 µg of sequencing grade trypsin (Promega) for 16 h at 37°C. Potential N-glycan chains were trimmed by the addition of 5 U of PNGase F for 1 h at 37°C followed by an additional step of trypsin digestion with 1 µg of trypsin for 3 h at 37°C. The SDC was removed by precipitation with 1% formic acid and centrifugation. Digested samples were cleaned up on Sep-Pak C18 cartridges (Waters) and dried in a vacuum centrifuge. Samples were reconstituted with 200 µL of 0.1% formic acid/4% acetonitrile. Sample concentrations were normalized based on the absorbance at 205 nm measured with a nanodrop (Thermo Scientific). Samples were spiked with the PQ500 kit (Biognosys), which consists of a mixture of 804 stable isotopes labeled peptides of known concentration. For reproducibility experiments, a pool of all samples was made and spiked with the PQ500.

### LC-MS data acquisition

The samples were analyzed by nano-UHPLC (nanoElute, Bruker Daltonics) coupled to a tims-TOF Pro mass spectrometer (Bruker Daltonics). Samples (4µl, 0.042 ng/µl) were directly injected onto a pulled emitter column (250 mm × 75µm, 1.6 µm, C18; IonOptiks) which was heated to 50°C in a column oven. Mobile phases consisted of 0.1% (v/v) formic acid in water (phase A) and acetonitrile (phase B). Samples were separated on a 100 min stepped gradient ranging from 2-30% B at a flow rate of 400 nl/min. The nano-UHPLC was coupled to a tims-TOF Pro instrument (Bruker Daltonics) operated in prm-PASEF mode or dia-PASEF mode. The prm-PASEF method was defined with a range of mobility values of 0.6-1.6 1/K0, a TIMS accumulation time fixed at 50 ms while the ion mobility separation was fixed to 100 ms. The time and mobility scheduled acquisition boxes were set with 2 min of tolerance on retention time and 0.05 1/K0 on the ion mobility.

The g-dia-PASEF method is based on dia-PASEF with 32 isolation windows of 26 m/z width, including a margin of 0.5 m/z. Isolation windows were associated with ion mobility windows of 0.3 1/K0 to cover as close as possible the peptide-ions distribution on both m/z and mobility dimensions. The TIMS accumulation and separation were both set at 100 ms.

### Data processing

All MS data were processed with Skyline daily^11^ and extracted fragment ion chromatograms (XICs) were extracted with a TOF resolution tolerance set to 60,000. For the g-dia-PASEF method, ion mobility data were filtered with tolerance windows of 0.05 1/K0 centered on the experimental mobility values and only fragment ions of the y-series were allowed, avoiding b-series crosstalk between the endogeneous and the isotope-labeled standard peptides. For g-dia-PASEF data, retention time prediction was performed using iRT peptides and using Biognosys spectral library values. The predicted retention time windows were set to 20 min. g-dia-PASEF transitions were manually curated to remove interfering transitions and correct wrong peak picking or peak integration boundaries.

The manually curated data were further processed using R (R-project.org). Peptides with incomplete measurement in all the samples were removed and the significantly regulated peptides were identified using a paired t-test corrected for multiple testing^12^. Peptides that showed an adjusted p-value < 0.01 were considered significantly regulated. The significantly regulated peptides were mapped back to the related protein.

### Bioinformatic meta-analysis

We have set up a meta-analysis and used it as previously described^13^. Briefly, we integrated all the individual CEL files from selected data sets profiled on HG-U133 plus 2.0 (Affymetrix, Santa Clara, CA, USA), retrieved from GEO (GSE14333, GSE17538, GSE21510, GSE8671, GSE9254, GSE20916, GSE10714, GSE15960, GSE4183, and GSE10961) and corresponding to different studies into one single global analysis covering expression data on 829 patients. The suitability of potential markers to discriminate between CRC and normal colorectal samples was assessed by ROC curves as previously described^13^.

## Results

### prm-PASEF and dia-PASEF acquisition methods design and performance

The PQ500 reference peptide kit consisted of 804 isotope-labeled peptides mixed at known concentrations and covering 579 blood proteins. Some strongly hydrophilic peptides did not elute reproducibly and thus were removed from the prm-PASEF method (supplemental table 1). The prm-PASEF method allowed the detection of 782 synthetic peptides spiked into the plasma samples, covering a total of 565 proteins with a 100 min chromatography gradient. Including the corresponding unlabeled peptides from the endogenous proteins, this represents a total of 1564 precursor ions analyzed with targeted acquisition windows of 2 min and ion mobility windows of 0.05 1/K0.

For the g-dia-PASEF method, variations in the retention times were not a problem as the method did not require scheduled acquisition. However, few peptides were not detected because they were outside of the m/z or ion mobility scanning range of the method (supplemental table 1). To ensure that no peptide was missing due to inaccurate retention time prediction, we manually re-analyzed all missing peptides without retention time filtering. We finally detected 756 internal standard peptides covering 549 proteins with the g-dia-PASEF method (supplemental table 2).

A PASEF event consisted of the accumulation of the incoming ions from the source into the first TIMS cell, the ion mobility separation of the peptide-ions in the second TIMS cell followed by the MS/MS analysis in the Q-TOF section of the instrument. The prm-PASEF MS cycle consisted of a variable number of PASEF events designed to acquire all the targeted peptides across the chromatographic separation. A prm-PASEF method is scheduled in a way that targets peptide-ions with a similar retention time but non-overlapping ion mobility values can be processed from the same ion mobility separation. If the ion mobility values overlap with another ion mobility, an extra PASEF event is added to the cycle time to address the situation. We recorded a maximum of 30 prm-PASEF events per MS cycle (figure 2A) which correspond to an MS cycle of approximately 3s. A maximum average of 6.8 precursor-ions targeted per prm-PASEF event in an MS cycle was measured (figure 2B). Despite the high density of targeted precursor-ions, it was possible to maintain a controlled cycle time with the prm-PASEF method and to acquire the peptides with a median number of 14 data points per peak and a minimum of 6 points (figure 2C) ensuring a proper quantification performance as illustrated by a median coefficient of variation of 3% (figure 2D).

**Figure 2.**
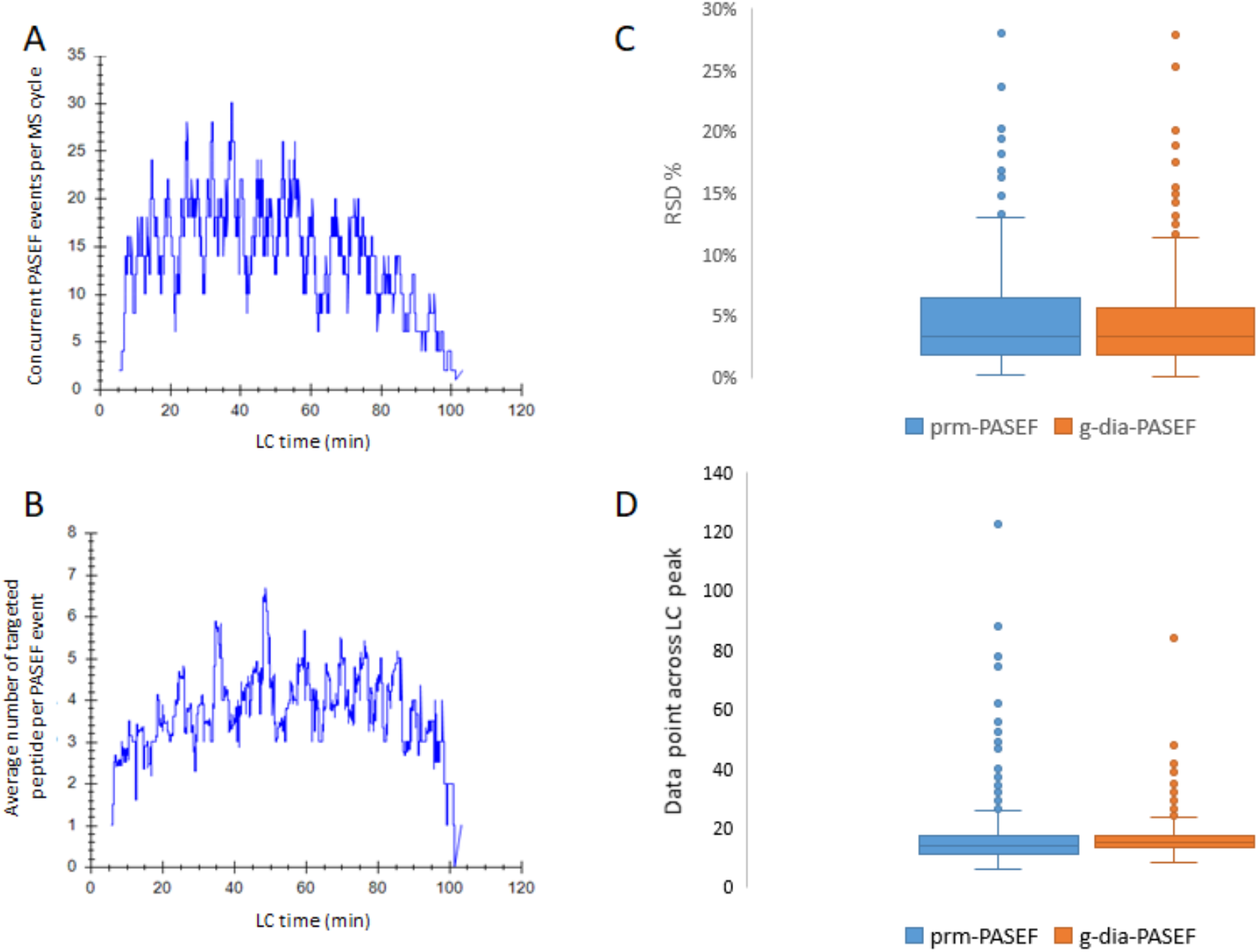
A) Number of PASEF events (100ms) per MS cycle across the chromatography separation, with the prm-PASEF method. B) Averaged number of PASEF events per MS cycle during the prm-PASEF acquisition C) Number of data points per LC peak profile. D) Relative standard deviation of the endogenous to heavy ratios of the prm-PASEF and g-dia-PASEF analysis a pooled plasma sample (n=3).

We observed that both prm-PASEF and g-dia-PASEF methods performed with similar reproducibility and a median relative standard deviation (RSD) of 3%. This value was calculated for each endogenous/heavy peptide pair detected in the three replicates. Because the MS cycle time was constant with g-dia-PASEF (1.8s) a sharper dispersion of the number of data points per peak was expected and the actual variation reflected the actual chromatographic peak width of the different peptides. prm-PASEF data were processed with the Skyline software and area ratios of the fragment-ion chromatograms of the endogenous and heavy-isotope-labeled standard as well as their respective dot-product similarity scores were exported. Data points associated with a dot product score below 0.98 were filtered out. We further manually inspected the chromatograms of the 10% lowest abundant peptides and eliminated peptides that were detected with poor signal quality. Figure 3A shows the data completeness distribution across the entire dataset showing a distribution where peptides are mostly detected in all or none samples for both methods. This distribution can be expected for a targeted acquisition method (i.e. PRM, prm-PASEF) or a targeted data processing (i.e. DIA, dia-PASEF) which are mostly driven by limits of detection. The risk of missing a peptide during the acquisition process is not as prevalent as with data-dependent acquisition (DDA). It was possible to estimate the actual concentration of the 572 endogenous peptides covering 378 proteins with the prm-PASEF method and this is 112 more peptides than with the g-dia-PASEF method (469 peptides quantified covering 308 proteins). This is principally due to the fact that PRM relies on narrow quadrupole isolation windows that improve the signal-to-noise ratio. The g-dia-PASEF method enabled the detection of 9 unique peptides that were typically peptides with variable retention times (figure 3B).

**Figure 3.**
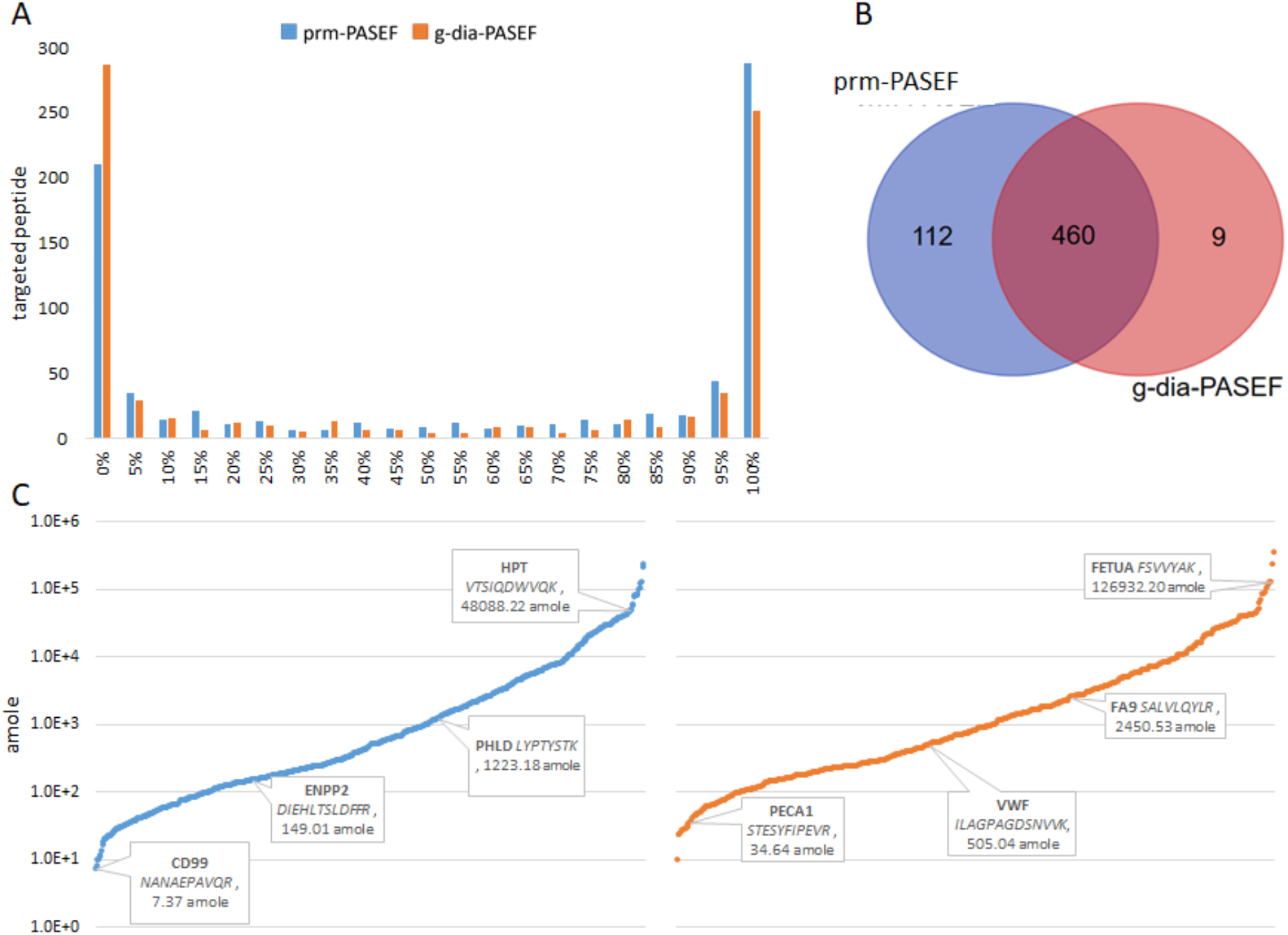
A) Histogram of the completeness for the prm-PASEF experiment. Bins represent the percentage of successful detection across the 20 samples. B) Peptides detection overlap between both methods. c) Dynamic range of the peptide detection. Peptides with different abundances are shown with the detected amounts.

We estimated the concentration of the endogenous peptides by single-point calibration with the spiked isotope-labeled standard (PQ500, Biognosys) (figure 3C). We calculated a peptide detection ranging from 7.4 (CD99) to 234232 (HEMO) amole injected onto the column for the prm-PASEF. The limit of detection and dynamic range is in line with our previous technical evaluation of the prm-PASEF method on serum and cell extract samples^9,14^. The prm-PASEF detected 27 low abundant peptides below 30 amoles injected whereas the dia-PASEF detected 8 peptides below that threshold. The correlation between the two methods for the overlapped quantification results was high (Supplementary figure 1, figure 4).

**Figure 4.**
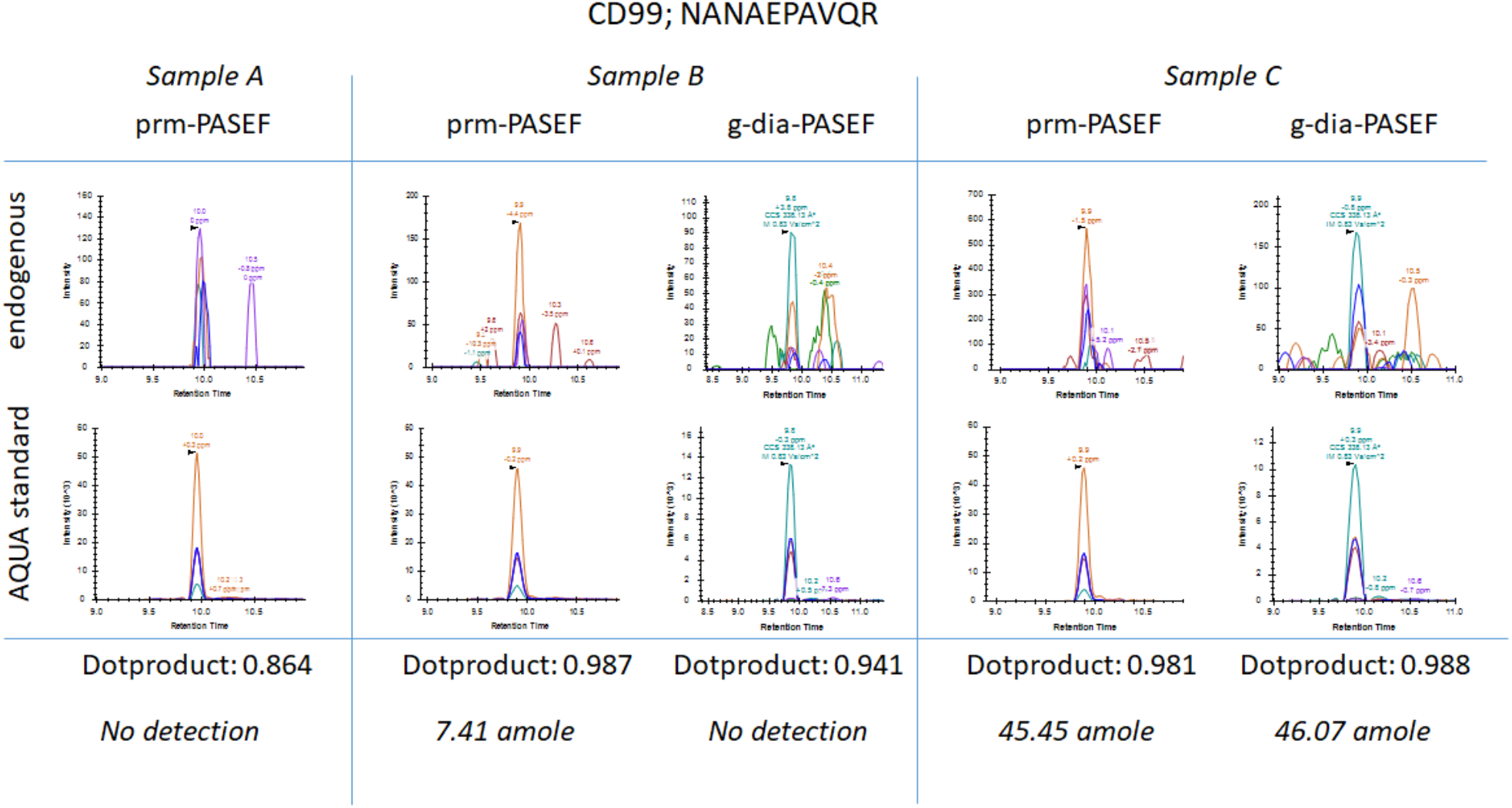
Extracted ion chromatograms of the peptides NANAEPAVQR (CD99) in prm-and g-dia-PASEF mode. Peptide identification was confirmed by similarity scoring (dot products) between the fragmentation pattern of the endogenous and the internal standard. Quantitation results are expressed in amole injected onto the column.

We finally compared the peptides and associated proteins detectable with label-free DIA data processing. It is indeed possible to process the DIA data without being restricted by the peptides included in the PQ500 kit and this approach detected (5773 peptides associated with 411 protein groups). Interestingly, we found that if the number of detected peptides is about ten times higher than targeted approaches, the number of detected protein groups stayed in the same range. In total 127 unique proteins were detected (supplemental figure 2) even if the label-free approach does not provide the same degree of identification and quantification confidence as SIL peptides based DIA, it still allows estimating the profiles of proteins not covered by the kit.

### Correlation of the prm-PASEF and g-diaPASEF data

The correlation between the two different measurement methods is high, with an R^2^ ranging between 0.92 to 0.97 for all samples. The main differences are the higher sensitivity of the prm-PASEF measurement, while the g-dia-PASEF method has a higher tolerance to retention time shifts.

The lower limit of quantification of the prm-PASEF method is based on the quadrupole peptide-ion isolation window which is at unit resolution. The gain in sensitivity comes at the cost of a more demanding method development. The prm-PASEF method requires experimental retention times and ion mobility values to set up the acquisition method. The acquisition method has to balance the number of peptides, the type of gradient, and MS measurement requirements to keep the number of data points per chromatographic peak acceptable for quantification.

For this analysis, we used a 25 cm column packed with 1.6 µm particles to increase the peak capacity while maintaining sharp elution profiles (*i*.*e*. 7.4s FWHM in average) on a 100 min gradient. The reproducibility of the retention was extremely important and required constant monitoring during the sample acquisition. Ion mobility values are susceptible to change with time and affect the synchronization of the TIMS with the quadrupole; we employed an automated recalibration of the ion mobility trap between each sample injection to alleviate this problem. The recently developed on-the-fly retention time correction might improve the implementation of prm-PASEF, allowing for narrower acquisition windows and thus shorter LC gradients^15^.

Conversely, the g-dia-PASEF method is more tolerant to retention time changes and does not require as much care and monitoring as prm-PASEF during the acquisition process. Since g-dia-PASEF is based on dia-PASEF it allows the *in silico* identification of peptides, which are not included in the peptide reference standard. Peptides without a reference standard can still be quantified between samples using label-free quantification methods (supplemental figure 2). The GO-analysis of the additionally identified proteins revealed that these are mainly belonging to the immunoglobulins and complement proteins (supplemental table 3).

One of the most time-consuming steps is the data processing and curation for prm-PASEF and g-dia-PASEF. The quality control of signal integration of 1564 precursor-ions is the main time-consuming step for both approaches. In our hands, Skyline was a practical software for visualization and correction of peak integration boundaries^11^. Skyline draws customizable and interactive graphics and tables that plot critical metrics such as retention times, XICs areas, mass error, and the dot-product similarity score. Graphics and tables are clickable and linked to the peptides XICs, allowing for a swift verification and correction of the outliers. We used the dot-product to filter true peptide signals from background noise. Additionally, each peak was manually reviewed, particularly for low abundance peptides. For these peptides background noise is particularly prone to give false-positive dot product scores. It is more likely to happen when one transition strongly dominates the others. We did not use Spectronaut (Biognosys) for the g-diaPASEF results processing because it tends to overestimate the peptides’ detection with this experimental design.

### Detection of differentially expressed proteins between CRC and control patients

To demonstrate the power of the prm-PASEF and g-dia-PASEF technique we applied the measurement to the plasma of 20 patients of the CRC cohort^16^. The patients of the two groups were age-matched (supplemental figure 3A) and the 10 CRC patients were equally distributed across the different disease stages (Supplemental figure 3B).

For the analysis of the prm-PASEF data only peptides, which were detected in all samples, were included. Of the 288 proteins which were detected in all samples, 16 were significantly regulated between the CRC patients and the control group (figure 5). While C4BPA, F13A, GELS, HGFA, IBP3, and IGF2 were downregulated in the CRC group versus the control group, A1AT, A2MG, C1QB, CD5L, CRP, CXCL7, HPT, PGRP2, and TRFE showed higher amounts in the CRC group (Figure 5). We used gene ontology term analysis to see if these proteins belong to a similar pathway. The majority of the proteins are associated with the activation of the immune system (supplemental figure 3). We used data obtained from different public tumor databases containing transcriptomics data obtained directly from tumor material^16^ to analyze if the genes were found to be associated with CRC in other studies (supplemental figure 4). The analysis showed that A2M, C1QB, C4BPA, CD5L, CRP, GSN (GELS_Human), ICAM2, IGF2, IGFBP3, PPBP, TF, and SerpinA1 mRNAs are upregulated in comparison to tissue samples of healthy control patients. Of those only the A2M, C1QBC4BPA, CD5L, CRPIGFBP3, TF, and SerpinA1 are specific to CRC tumor samples (supplemental figure 4). An analysis of the current literature revealed that A1AT^17^, ICAM2^18^, IGF2^19^, F13A^20^, CXCL7^21^, and SHBG^22^ have been previously found to be associated with colon cancer.

**Figure 5.**
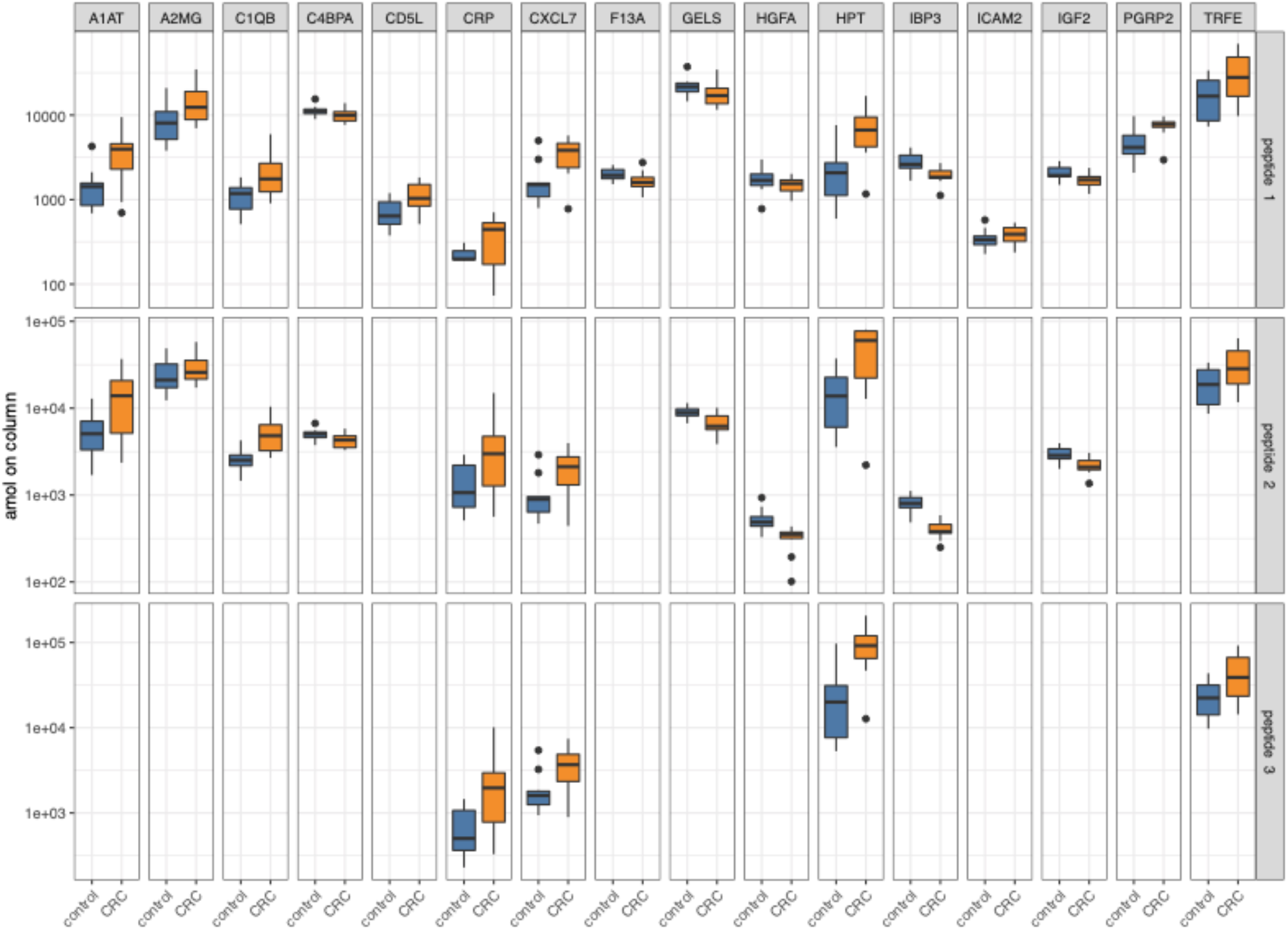
Boxplot for the proteins significantly regulated between the CRC and control patients. In the PQ500 standard, the number of peptides per protein varies. For the regulated proteins, the number of peptides per protein varies between one and three peptides. Each boxplot shows the distribution of the quantification separated by each peptide. CRC patients are shown in orange and controls in dark blue.

## Conclusions

Here, we describe the use of a highly complex isotope-labeled peptide standard in combination with prm-PASEF and dia-PASEF. The combination of the highly complex peptide reference standard, PQ500, allowed us to develop the new g-dia-PASEF technique, which is less complex to set up than the prm-PASEF method, while still allowing the absolute quantification of the plasma proteins.

This highly multiplexed analysis allowed us to measure and quantify 782 peptides in plasma samples with a high dynamic range of four orders of magnitude. The prm-PASEF method identifies the highest number of proteins in a plasma sample but requires a significant effort to set up the acquisition method. An alternative is the use of the g-dia-PASEF method, which combines the ease of setting up a dia-PASEF method with the possibility to quantify proteins to a reference standard. The use of a reference standard offers the important advantage of cross-experiment comparison which is essential for the use of biomarker measurements.

The careful analysis of each peptide’s XIC (extracted ion chromatogram) revealed that the automated software algorithms for peaks detection, integration, and identity validation are still very optimistic. A more stringent reevaluation of the results is necessary for the targeted measurement using prm-PASEF and g-dia-PASEF. This is particularly important for potential clinical applications.

Using the prm-PASEF and g-dia-PASEF methods for analyzing the CRC-cohort samples we showed that with an unbiased measurement of the 579 selected plasma proteins we are able to distinguish CRC-patients from the control group, by measuring a representative portion of the plasma proteome composition. The significantly regulated proteins contain proteins, which are associated with colon carcinoma or with immune responses, which are typical for CRC patients. For validation of these proteins as a biomarker more measurements in a larger cohort will be necessary, but the 16 proteins already underline the potential of the measurement as a diagnostic tool.

Based on this study it should be possible to further utilize this technique as a general measurement standard. This means that a patient sample can be measured using the PQ500 or a similar standard and the results can be directly compared to other patient samples measured using the same approach. The diagnostic potential would be extended by the measurement of 500 proteins and will allow the definition of protein panels as biomarkers of different diseases. A single plasma proteome measurement using the PQ500 standard with either prm-PASEF or g-diaPASEF could be the unified diagnostic tool for many different diseases.

## Acknowledgment

We thank the Fondation Cancer for the support with grant FC/2022/01 SOCS to EL. The authors would like to thank Dr Guy Berchem and Dr Keipes for their support and the IBBL for the overall setup of the sample collection, processing, and provision. We thank Komal Baig for the setup of the meta-analysis.

## Authors contribution

AL developed the method implementation. AL and FB performed the sample preparation and measurement. EK performed the analysis of the tumor expression data. EL provided the CRC samples. AL, GK, POS, and GD developed the methodology. AL wrote the first draft. AL and GD finalized the manuscript.

## Supplementary Material

**Supplementary figure 1.**
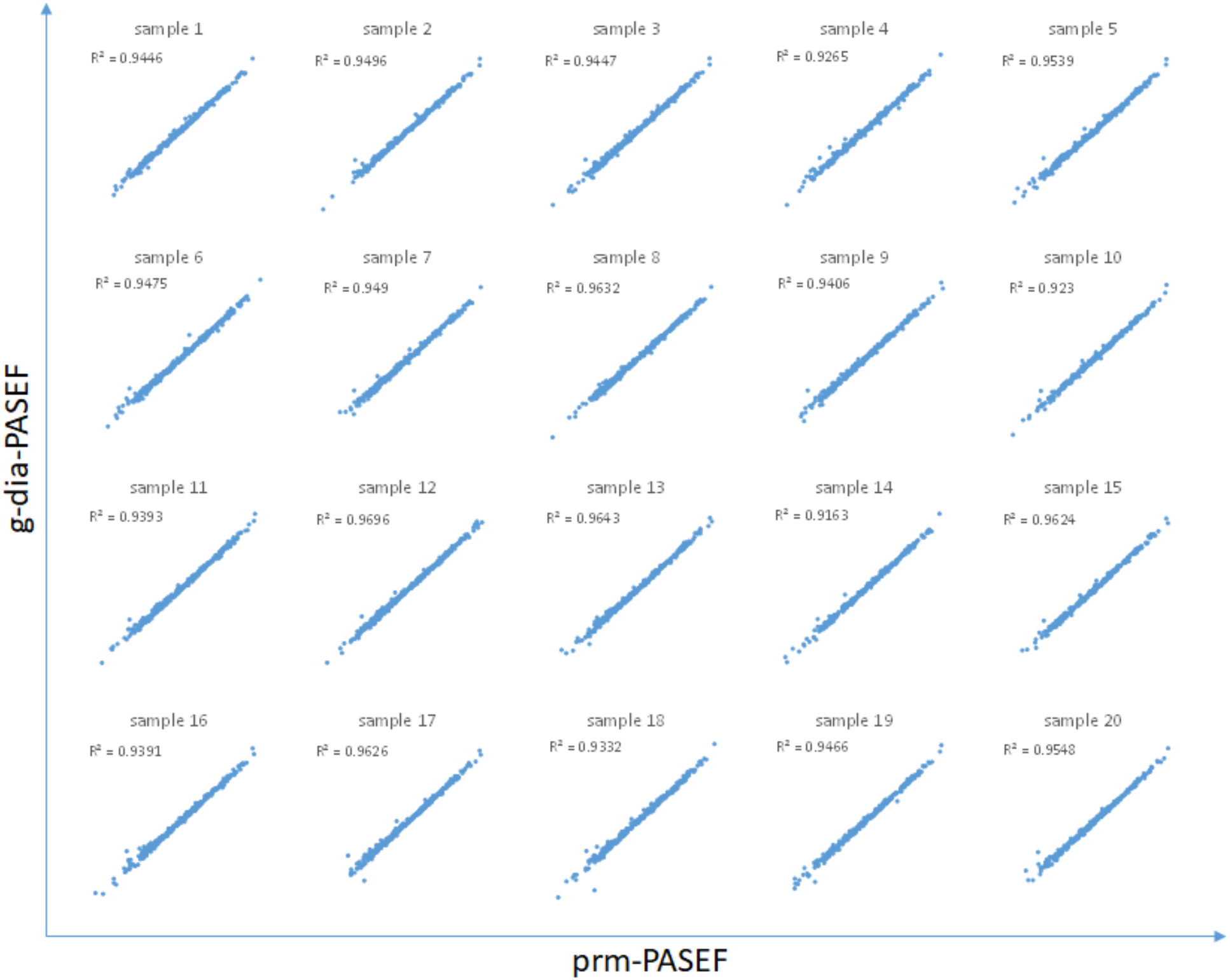
Correlation of the peptide quantification results between g-dia-PASEF and prm-PASEF acquisition. Axis are in logarithmic scale.

**Supplemental figure 2.**
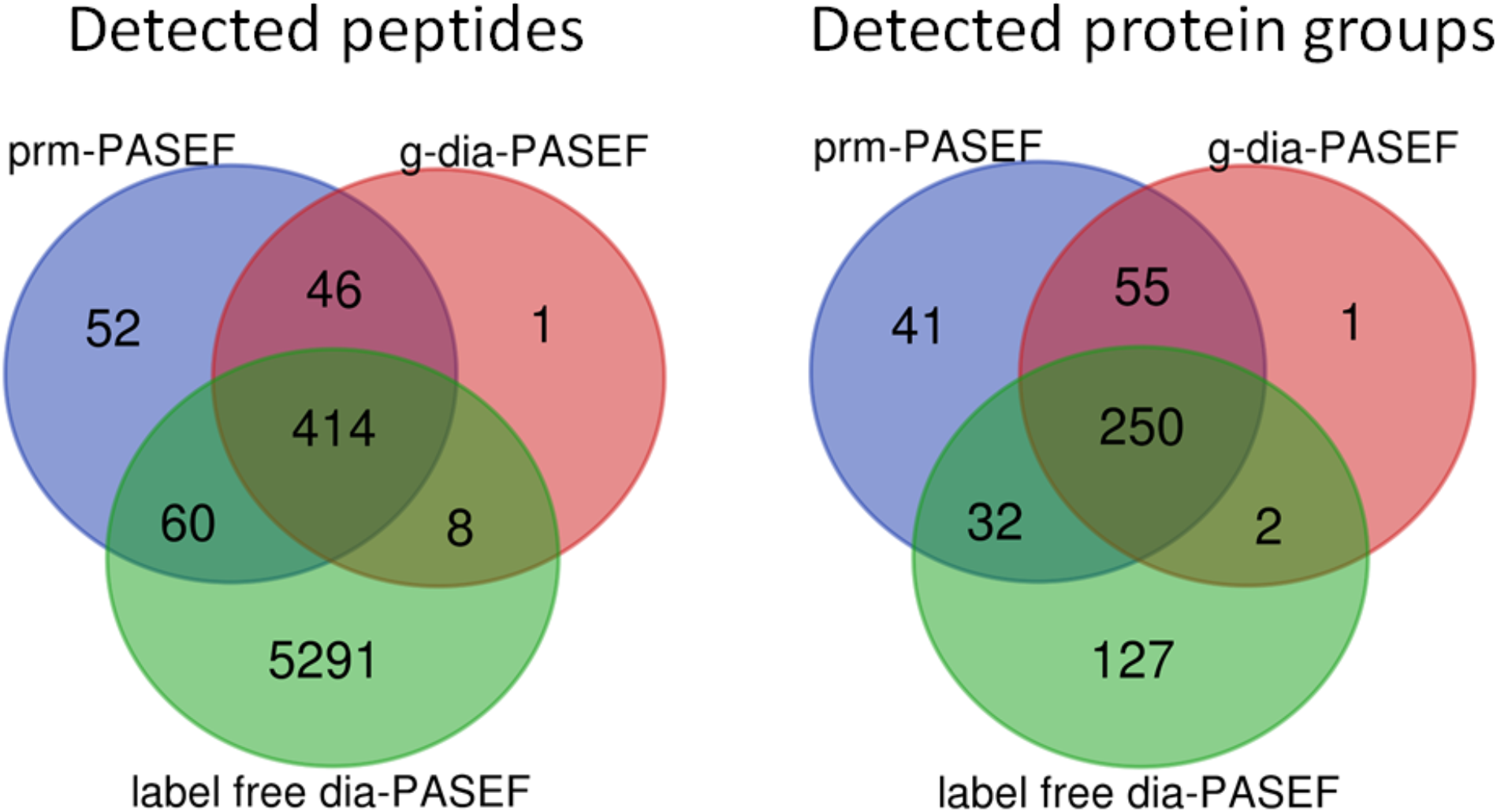
Peptides detection overlap between label-free dia-PASEF, g-dia-PASEF, and prm-PASEF methods. g-dia-PASEF and prm-PASEF methods detected 97 unique proteins. From the 127 unique proteins detected by label-free dia-PASEF, 96 were not covered by the PQ500 kit, 10 were detected with a peptide not covered or not detected from the PQ500 kit and finally, 21 corresponded to a very low signal that did not pass the identification threshold in g-dia-PASEF.

**Supplemental figure 3.**
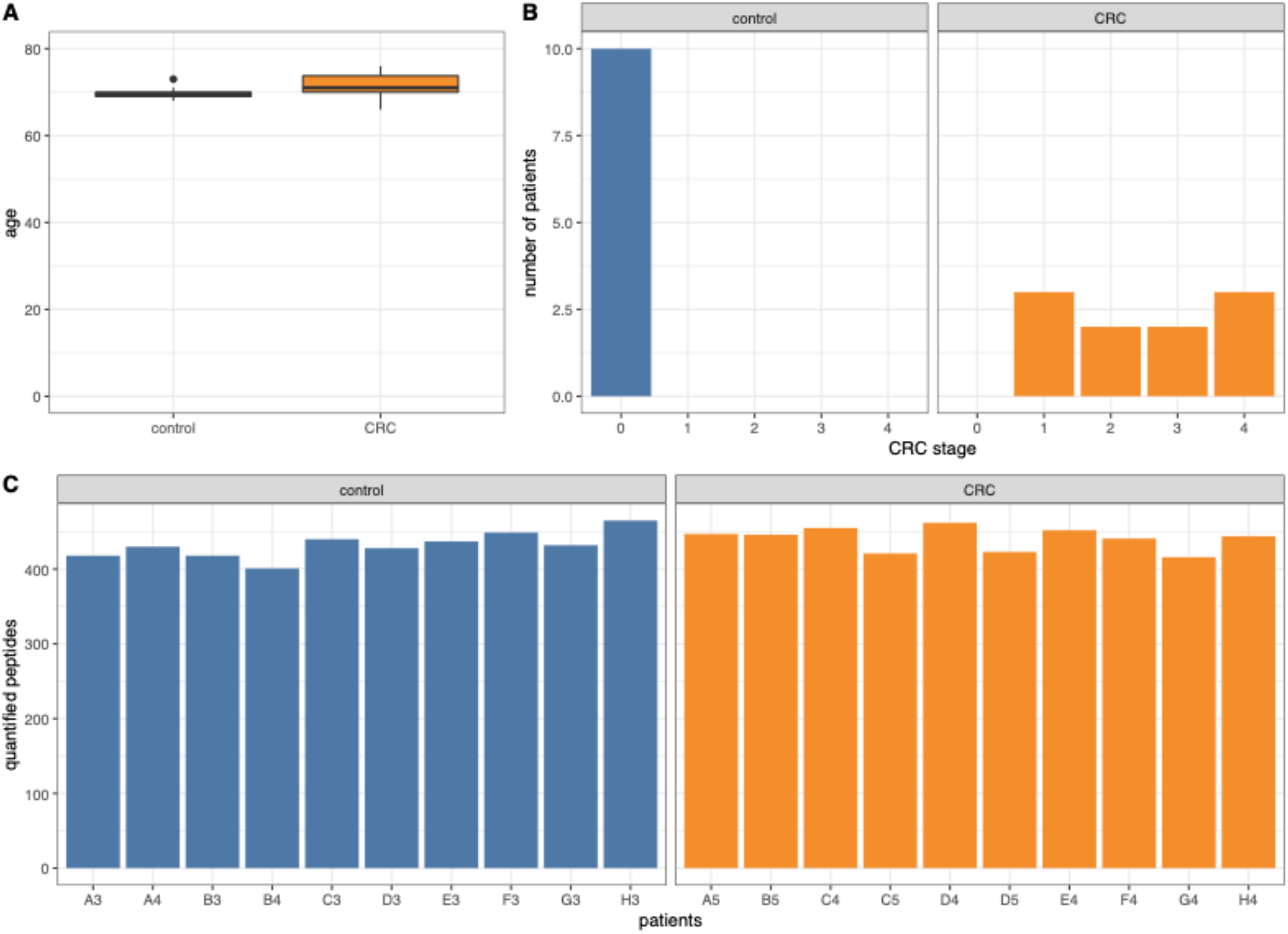
**A**. Age distribution of the 10 CRC and the 10 control group patients. **B**. Distribution of the CRC patients across the different CRC stages. The control samples from healthy individuals are listed as stage 0. **C**. Peptides identified by prm-PASEF in each of the patient samples.

**Supplemental figure 4.**
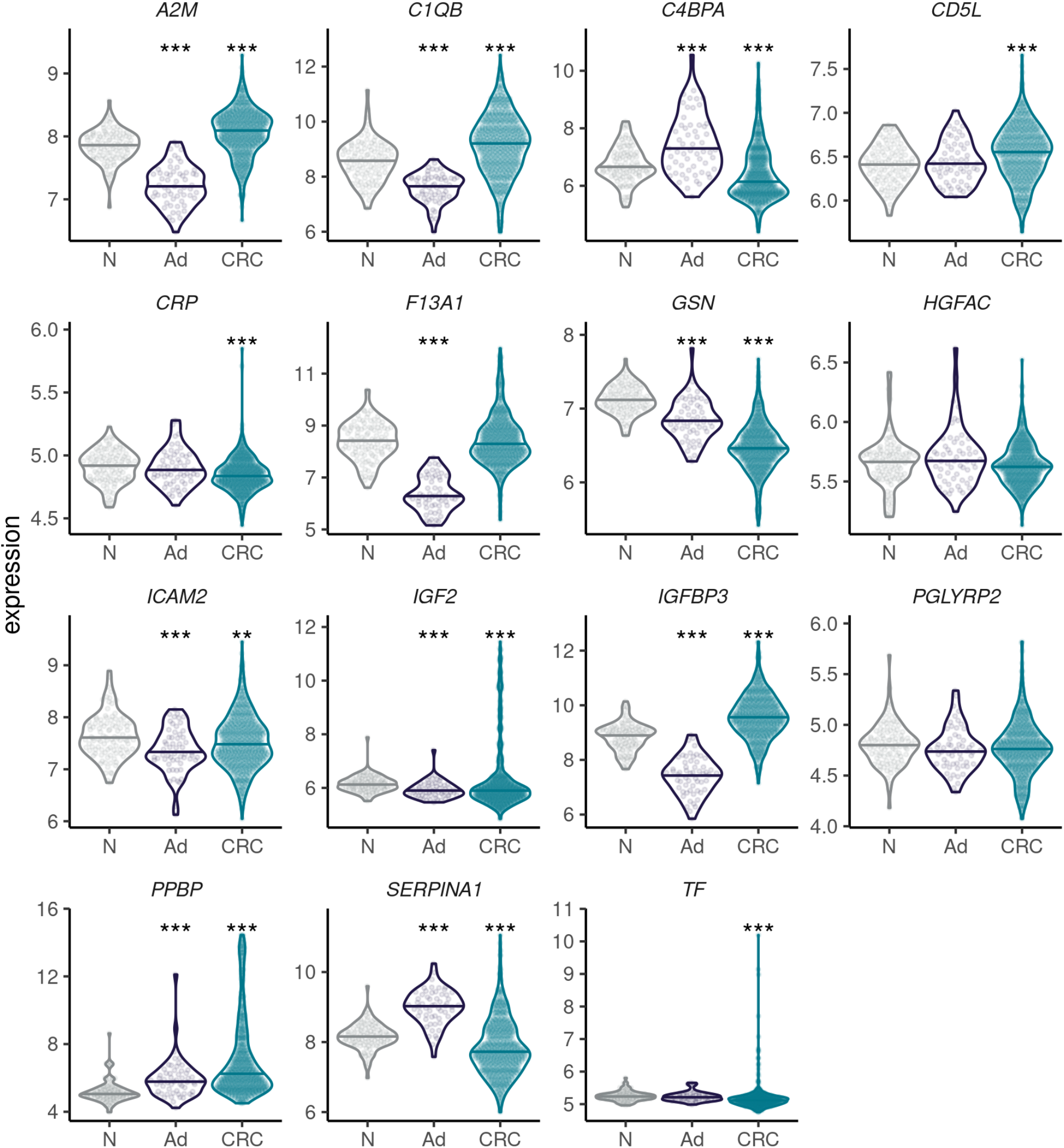
Expression of genes corresponding to the identified proteins in tissues of CRC patients. Beeswarm and violin plots showing the normalized expression values of the 15 genes in adenoma (n = 58) and CRC (n = 673) samples compared to healthy colorectal mucosa samples (n = 102) in a meta-dataset composed of different publicly available CRC datasets, totalizing 833 patients. Horizontal lines represent the median expression value (*** p < 0.001; ** p < 0.01; Kruskal-Wallis test followed by pairwise Wilcoxon-rank sum tests against the normal tissue sample conditions. p-values were corrected for multiple comparisons using Holm’s method).

**Supplementary table 1:**List of detectable SIL peptides with prm-and g-dia-PASEF.

→ see excel file.

**Supplemental table 2.**
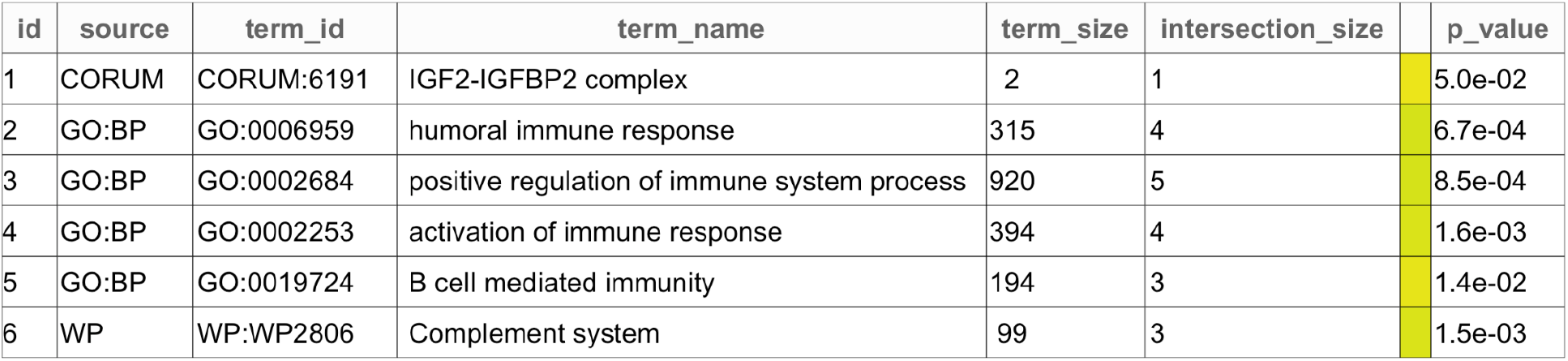
GO term analysis of the significantly regulated proteins.

**Supplemental table 3:** GO term analysis of the proteins identified in the label-free dia-PASEF analysis.

| id | source | term id | term name | term size | intersection size | p-value |
| --- | --- | --- | --- | --- | --- | --- |
| 1 | GO:CC | GO:1903561 | extracellular vesicle | 2132 | 67 | 9.5e-35 |
| 2 | GO:BP | GO:0006956 | complement activation | 128 | 22 | 7.9e-24 |
| 3 | GO:CC | GO:0019814 | immunoglobulin complex | 165 | 21 | 1.1e-20 |
| 4 | GO:MF | GO:0003823 | antigen binding | 166 | 19 | 4.0e-17 |
| 5 | GO:CC | GO:0042571 | immunoglobulin complex, circulating | 75 | 14 | 9.9e-16 |

