## Supplementary figures and images for "Highly multiplexed targeted plasma proteomics quantifies several hundred blood proteins in serum from colorectal carcinoma patients"

### supplementary figure 1

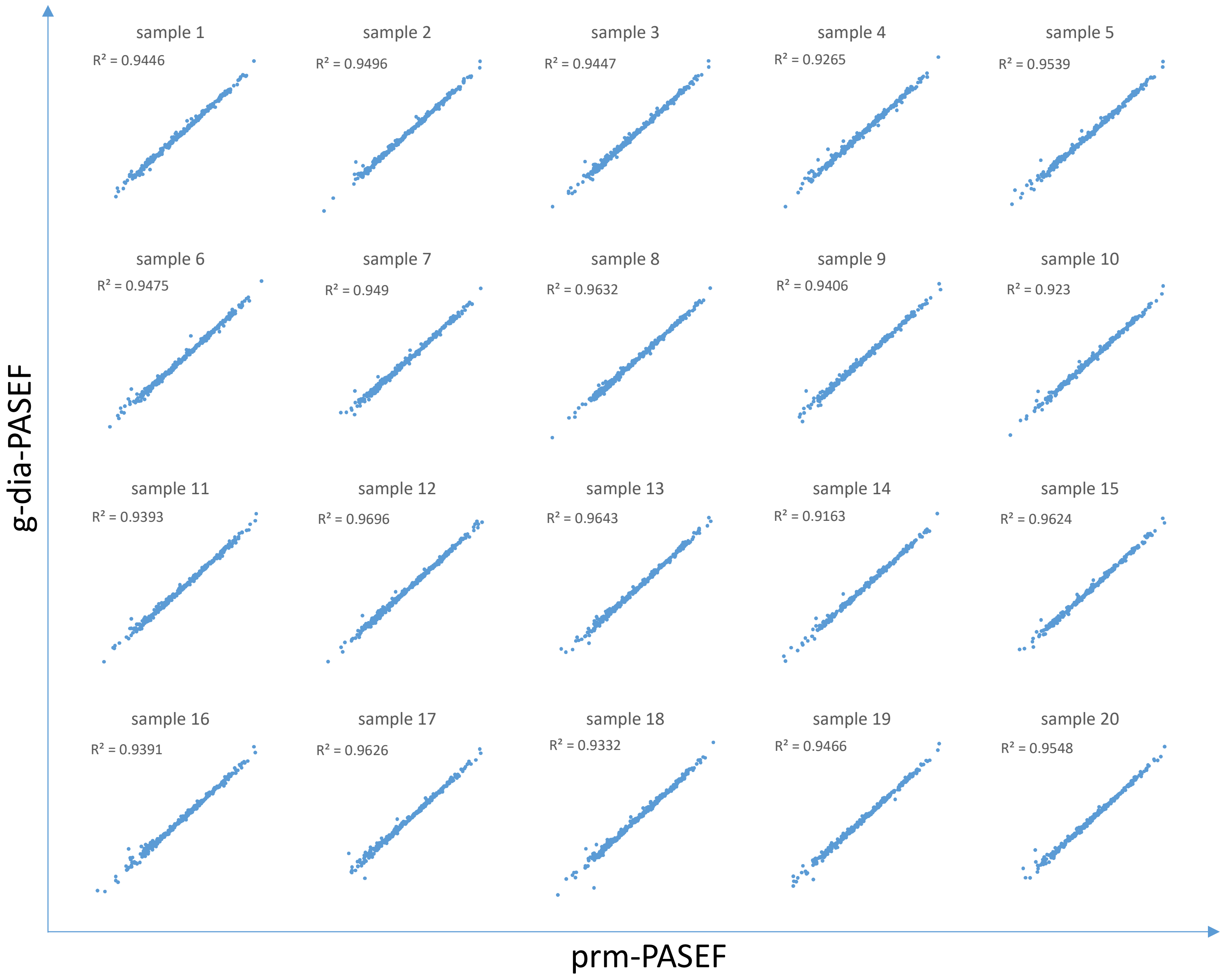

### supplementary figure 2

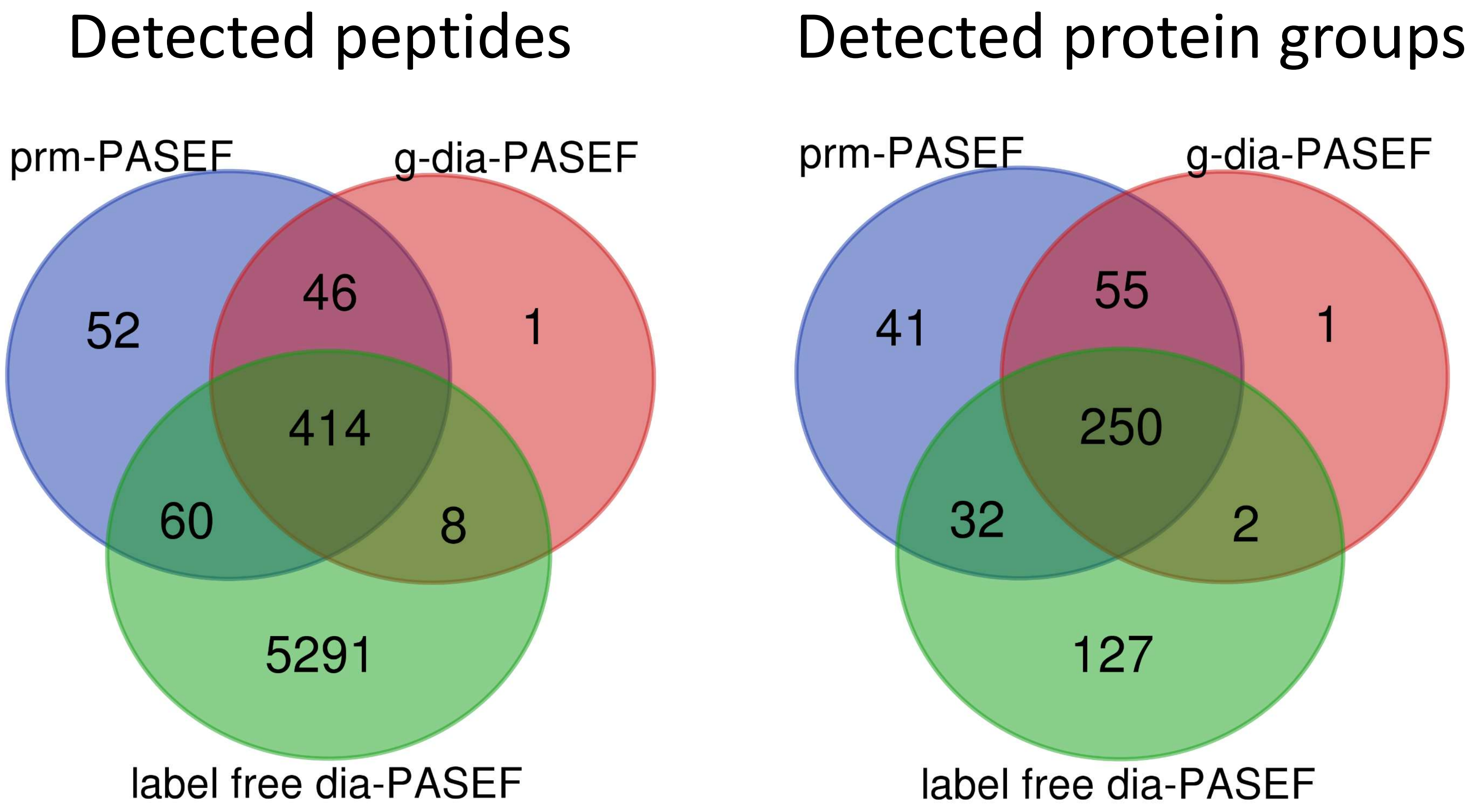

### supplementary figure 3

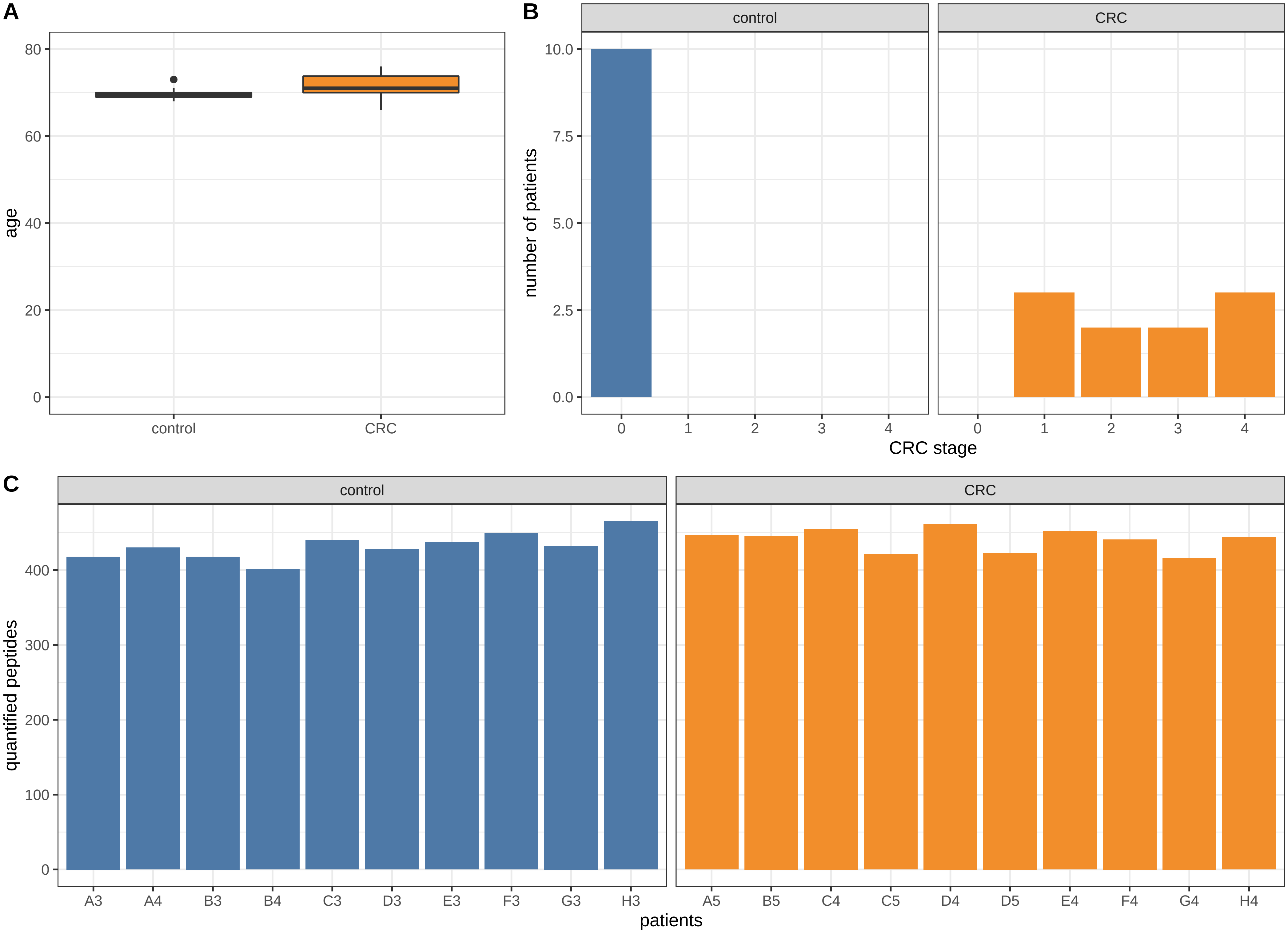

### supplementary figure 4

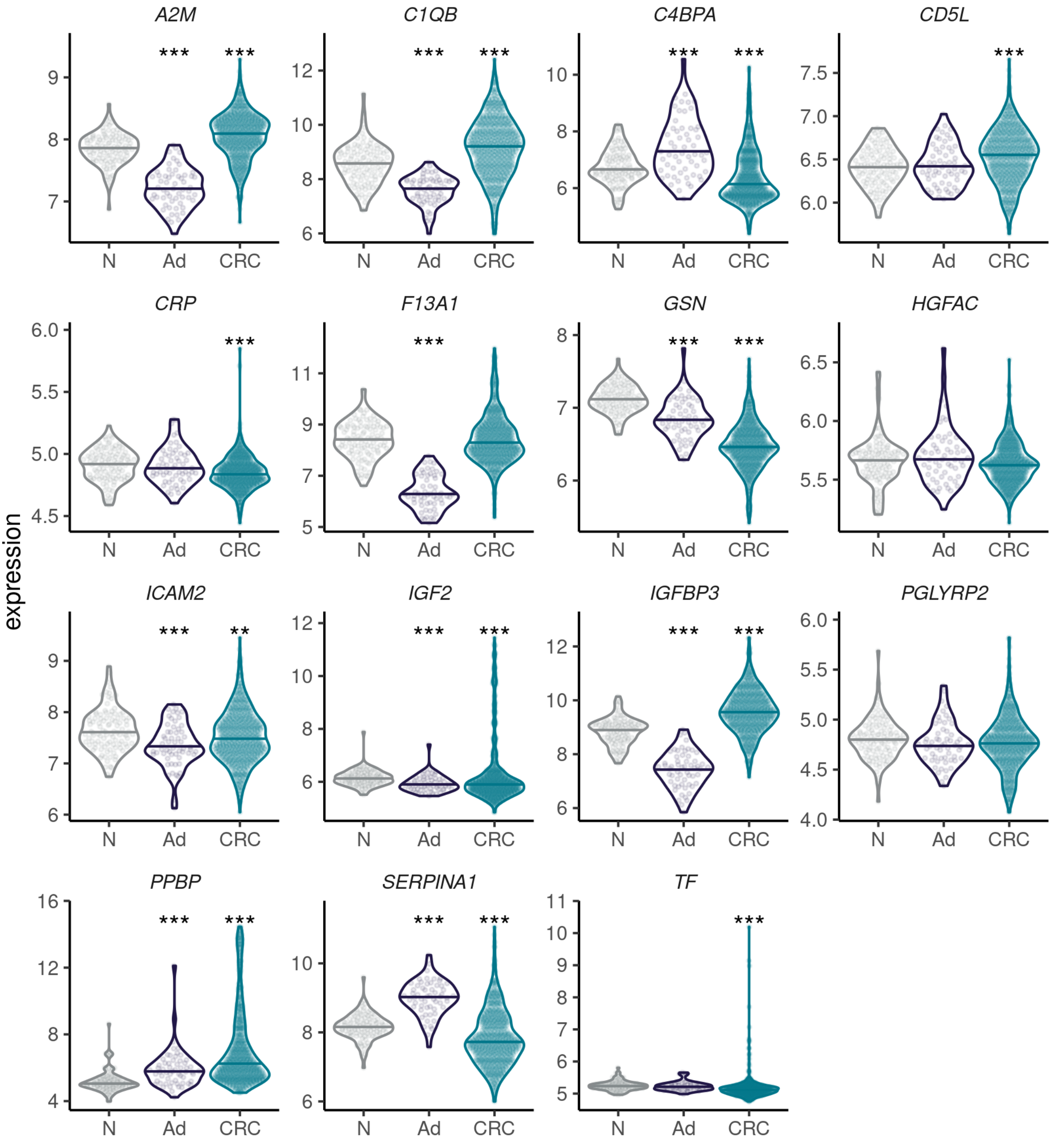
